# Models for infantile hypertrophic pyloric stenosis development in patients with esophageal atresia

**DOI:** 10.1101/625921

**Authors:** Chantal A. ten Kate, Rutger W.W. Brouwer, Yolande van Bever, Vera K. Martens, Tom Brands, Nicole W.G. van Beelen, Alice S. Brooks, Daphne Huigh, Bert J.F.M.M. Eussen, Wilfred F.J. van IJcken, Hanneke IJsselstijn, Dick Tibboel, Rene M.H. Wijnen, Annelies de Klein, Robert M.W. Hofstra, Erwin Brosens

**Author notes:** **Corresponding Author:** E. Brosens, PhD, Erasmus Medical Centre – Sophia Children’s Hospital, Room Ee-987e, P.O. Box 2040, 3000 CA Rotterdam.

## Abstract

Patients born with esophageal atresia (EA) have a 30 times higher prevalence of infantile hypertrophic pyloric stenosis (IHPS). This makes sense from a developmental perspective as both the esophagus and the pyloric sphincter are foregut derived structures. EA and IHPS are variable features in several (monogenetic) syndromes. This, and twin and familial studies, indicates a genetic component for both conditions as single entities. We hypothesized that genetic defects, disturbing foregut morphogenesis, are responsible for this combination of malformations. Non-genetic factors could also contribute, as mice exposed to Adriamycin develop EA and *in utero* diethylstilbestrol exposure is associated with EA.

We investigated the copy number profiles and protein coding variants of 15 patients with both EA and IHPS. As all parents were unaffected, we first considered dominant *(de novo)* or recessive inheritance models but could not identify putatively deleterious mutations or recessive variants. We did identify inherited variants in genes either known to be involved in EA or IHPS or important in foregut morphogenesis in all patients. Unfortunately, variant burden analysis did not show a significant difference with unaffected controls. However, the IHPS associated risk SNP rs1933683 had a significantly higher incidence (OR 3.29, p=0.009).

Although the genetic variation in likely candidate genes as well as the predisposing locus near *BARX1* (rs1933683) suggest a genetic component, it does not fully explain the abnormalities seen in these patients. Therefore, we hypothesize that a combination of high impact genetic, mechanical and environmental factors together can shift the balance to abnormal development.

**Summary statement:** Instead of one affected gene, the higher incidence of IHPS in EA patients is more likely the result of multiple (epi)genetic and environmental factors together shifting the balance to disease development.

## INTRODUCTION

Esophageal atresia (EA) is a rare congenital malformation caused by a faulty development of the foregut which leads to a discontinuity of the esophagus. It occurs in about 2.5 cases per 10,000 births within Europe (Pedersen et al., 2012, Oddsberg et al., 2012) and over three-quarters of patients present with a tracheoesophageal fistula (TEF) (Pedersen et al., 2012, Macchini et al., 2017). EA is considered – etiologically as well as phenotypically – a highly heterogeneous condition (Brosens et al., 2014). It can present either as an isolated defect but is often seen in combination with other malformations. Frequently, these malformations are part of the VACTERL (Vertebral, Anorectal, Cardiac, Tracheoesophageal, Renal or urinary tract of Limb malformations) association. VACTERL association is a diagnosis of exclusion in which three or more features of the VACTERL spectrum are present and a known genetic syndrome is not identified (Solomon et al., 2012). However, clustering of one or more of these features with additional specific associated malformations could also be the results of a shared genetic etiology.

One of the more prevalent, but less well-known, associated malformations is Infantile Hypertrophic Pyloric Stenosis (IHPS) (Rollins et al., 1989). In contrast to EA, IHPS is often considered an acquired disorder. The pyloric muscle hypertrophies in the first weeks of life, causing a narrowing of the pyloric channel (Panteli, 2009). Healthy-born infants present at week 3 to 6 of life with projectile postprandial vomiting. They need surgery where the upper layer of the circular smooth muscle of the pylorus will be incised, to release the passage from the stomach to the intestine again.

Previously, we described a 30 times higher prevalence (7.5%) of IHPS in patients with EA compared to the normal population (0.25%) (van Beelen et al., 2014). This increased prevalence has been reported in other retrospective studies (3.3-13%) as well (Palacios M.E.C. et al., 2014, Deurloo et al., 2002). The clinical presentation of IHPS seen in patients with EA/IHPS is not different from patients with isolated IHPS. However, the diagnosis of IHPS is more difficult and often delayed in patients with EA. Relatively common complications after EA repair, such as stenosis of the anastomosis, can protect against reflux and lead to just regurgitation. By the time these patients start vomiting, there must be massive gastroesophageal reflux.

The presentation of both EA and IHPS makes sense from a developmental perspective as the esophagus and the pyloric sphincter are both foregut derived structures. Organ specification during embryonic development is under tight spatiotemporal control of specific growth factors, transcription factors and signaling cascades (Li et al., 2009, Jacobs and Que, 2013). Disturbances in these pathways could impact proper development. In mice, the esophagus is specified from the foregut tube between embryonic day E9.5 and E11.5. In humans, the esophagus, as well as the stomach, starts developing from the fourth week after conception onwards. The stomach turns around its anterior-posterior axis during embryonic development (Cetin et al., 2006). The developing pylorus can be visualized with immunostaining at week six after gestation and differentiates during fetal life (Koyuncu et al., 2009).

Environmental (Zwink et al., 2016, Felix et al., 2008, Feng et al., 2016, Markel et al., 2015, Krogh et al., 2012, Sorensen et al., 2002) and genetic contributions (Peeters et al., 2012, Brosens et al., 2014, Solomon et al., 2012) have been described for both EA and IHPS as single entities or in combination with other anatomical malformations. It has been suggested that *in utero* exposure to diethylstilbestrol (DES) is associated with the development of EA (Felix et al., 2007a). Moreover, both malformations are variable features in specific and often phenotypically overlapping genetic syndromes (Table 1). The presence of both conditions as variable features in the phenotypical spectrum of known genetic syndromes is indicative of a genetic background for EA and IHPS. More evidence for a genetic contribution can be deduced from twin studies and animal models (de Jong et al., 2010). The concordance rates in monozygotic twins compared to dizygotic twins is higher for EA (Veenma et al., 2012) and IHPS (Krogh et al., 2010) as single entities. Also, the recurrence risk is elevated for siblings and offspring of affected individuals with EA in combination with other associated anomalies (Robert et al., 1993, Van Staey et al., 1984, Warren et al., 1979, McMullen et al., 1996). In contrast, the recurrence risk for isolated EA is low (Schulz et al., 2012) and moderate for IHPS (Krogh et al., 2010, Elinoff et al., 2005). In contrast to EA, there has been reported a male predominance for IHPS (4:1) (MacMahon, 2006). There have been risk loci associated to IHPS (Everett and Chung, 2013, Feenstra et al., 2012, Feenstra et al., 2013, Svenningsson et al., 2012, Fadista et al., 2019). To date, no risk loci have been described for EA.

**Table 1.**
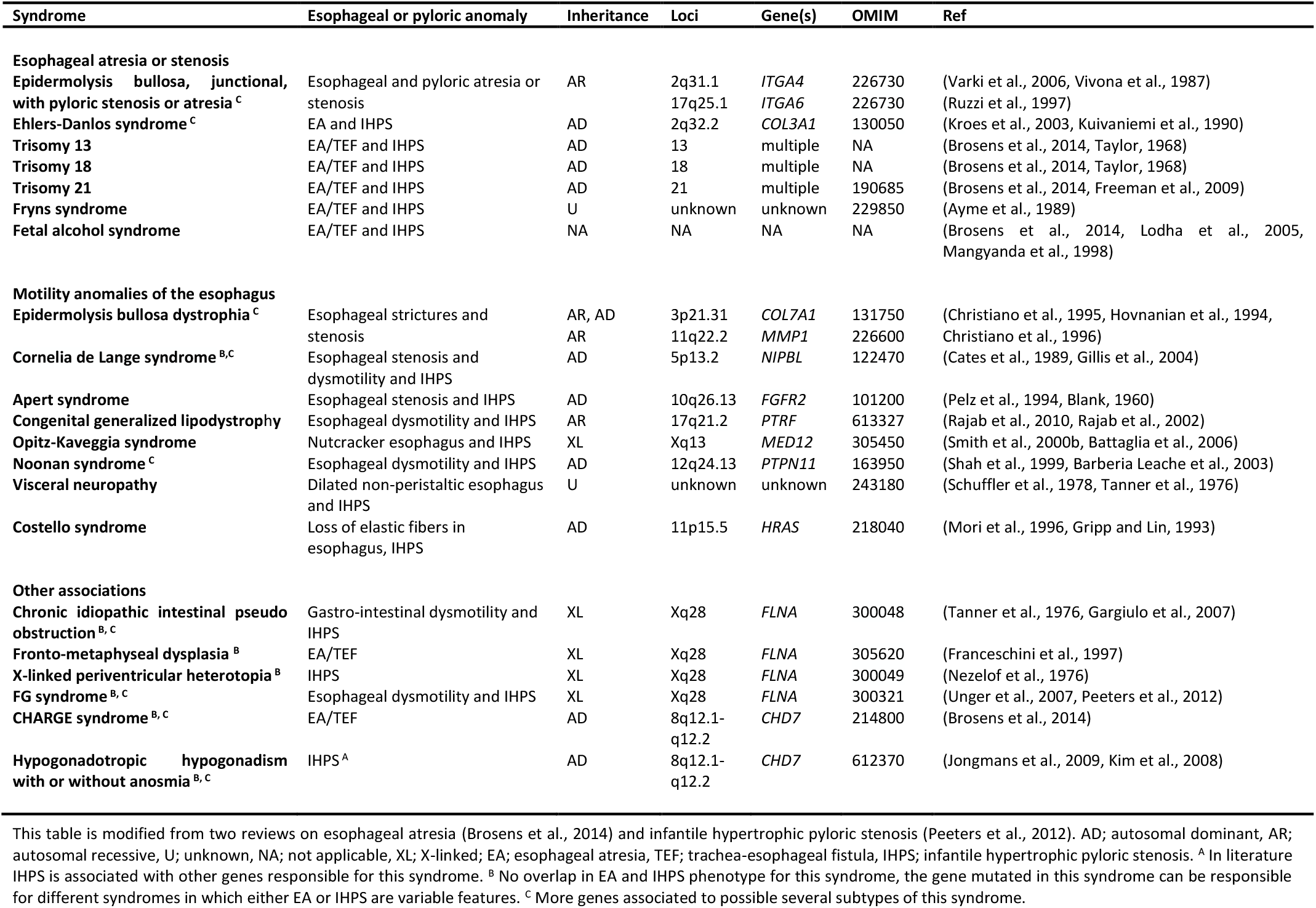
Genetic syndromes and mutated genes with tracheoesophageal and pyloric anomalies as variable features

Considering the increased prevalence of IHPS in patients with EA, their common developmental origin and previous evidence in genetic studies, we hypothesized that Copy Number Variants (CNVs) or other protein coding alterations affecting one specific gene, or genetic disturbances in more genes all important for foregut morphogenesis are responsible for the higher incidence of IHPS in patients with EA.

## RESULTS

### Patient cohort

In total, 27 out of 664 patients (4.1%) born with EA between 1970-2017, developed IHPS. Twenty patients have been described previously (van Beelen et al., 2014). Parental informed consent for whole exome sequencing (WES) was obtained for 15 patients. Several phenotypical characteristics stood out in this EA/IHPS cohort: a sacral dimple was present in seven patients (25.9%), anomalies of the vertebrae or ribs in eight patients (29.7%) and genitourinary anomalies in six patients (22.2%) of which two patients (7.4%) had hypospadias. Four patients (14.8%) had three or more anomalies within the VACTERL spectrum (Solomon, 2011). A full phenotypical description of the 27 EA/IHPS patients is given in Table 2.

**Table 2.**
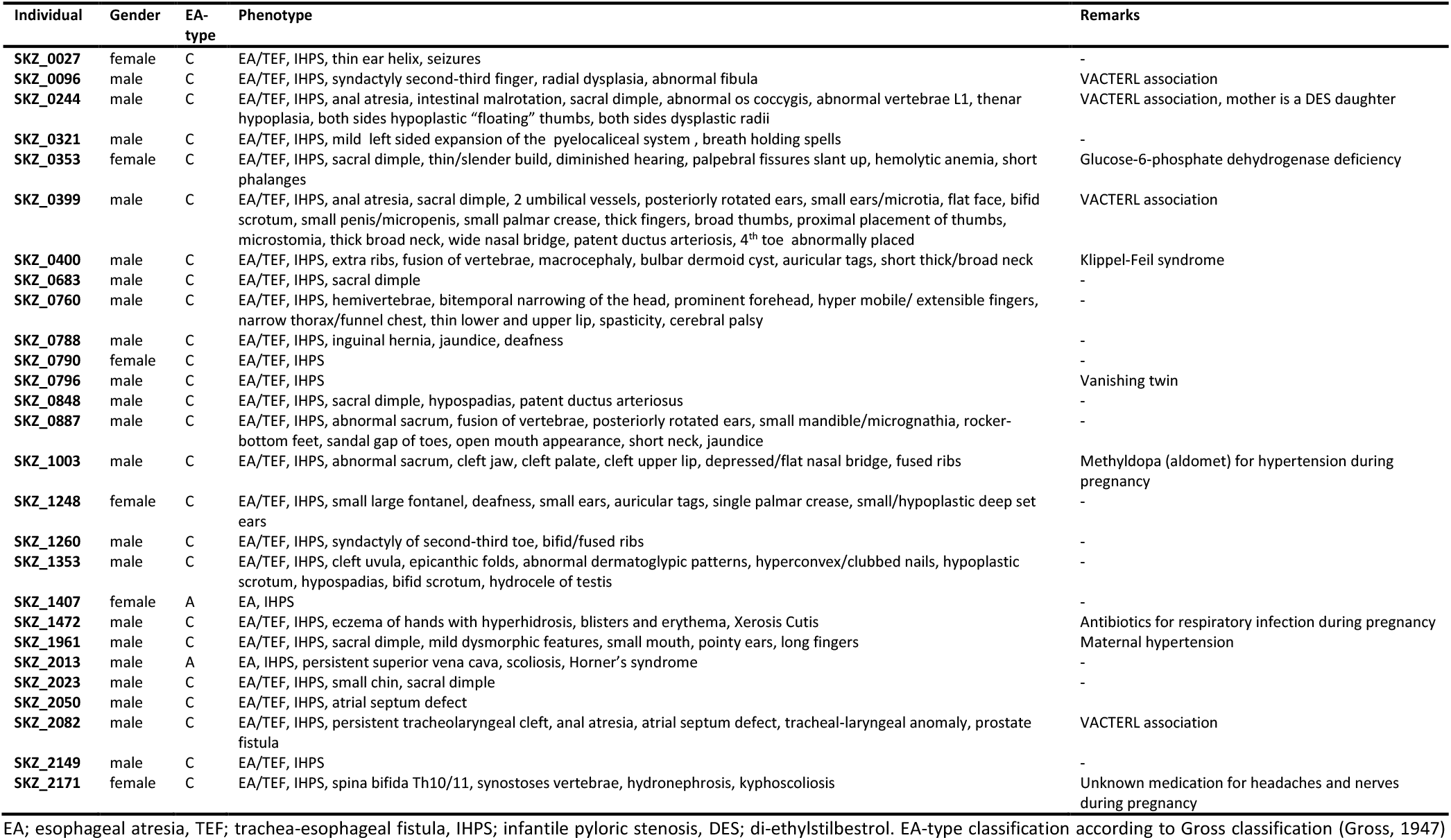
Phenotype description

### Copy Number analysis

Our previous study described rare CNVs and their inheritance pattern in patients with EA (Brosens et al., 2016b), seventeen EA/IHPS patients were included in this previous study. None of the six large CNVs identified were *de novo,* all were inherited from one of the unaffected parents. Patient SKZ_400 had a paternal inherited rare gain of chromosomal region 11q15. Patient SKZ_0887 had maternal inherited putative deleterious gains on Xq26.1 and Xp22.33. Patient SKZ_1003 had a maternal inherited loss of chromosomal region 17q11 and patient SKZ_1248 maternal inherited rare gains in chromosomal regions 4q35 and 5p15.1. Additional exon-level CN-profiling using the normalized coverage profiles (Amarasinghe et al., 2013) of the exome sequencing data confirmed the presence of the CNV seen with SNP-array. All CN profiles of main EA and IHPS disease genes (Brosens et al., 2014, Peeters et al., 2012) were normal. There were no overlapping rare CNVs in this patient cohort. All rare CNVs, classified as VUS or (likely) deleterious are described in Table S1.

### Exome sequence analysis

Sequencing resulted in at least 5 Giga-bases of raw sequence data with an average coverage of 70X and 90% of target bases covered over 20X. Quality of the sequence data is listed in Table S2. As none of the parents of the 15 investigated patients were affected we first considered dominant *de novo* and recessive modes of inheritance.

We could not identify *de novo* pathogenic variation in main EA and IHPS disease genes (Brosens et al., 2014, Peeters et al., 2012). Subsequently, we searched for possible *de novo* mutations exome wide. For this, we focused on putative deleterious ultra-rare protein coding or splice site variants (n=100) (Bennett et al., 2017). Variations were considered ultra-rare when they were absent in the gnomAD dataset (123,136 whole exomes and 15,496 whole genomes) (http://gnomad.broadinstitute.org/) (Lek et al., 2016). Twenty-five variants proved to be sequencing artifacts. Furthermore, we could not confirm the segregation of 15 mutations due to lack of parental DNA. We determined the segregation of all ultra-rare variants predicted to be of unknown significance (VUS, n=37) or (likely) deleterious (n=23). All putative deleterious variants tested proved to be inherited from one of the unaffected parents.

Considering a recessive mode of inheritance, we searched for genes with homozygous or compound heterozygous variants. Six variants in three genes *(FLNC, ATP6V0A1* and *FAM46A)* fitted a putative compound heterozygous model, two genes *(KCNN3* and *VDAC3)* had homozygous variants and two genes *(MID2* and *SH3KBP1)* had variants on chromosome X in a male patient. All variants were predicted to be likely deleterious or VUS and intolerant to missense variants (Z-score ≥3) or loss of function variants (PLI or PLIrec ≥0.9). With segregation analysis, we could confirm the compound heterozygous mode of inheritance of the variant in the *FAM46A* gene in patient SKZ_2023 and the maternally inherited X-linked variant in the *SK3KBP1* gene in patient SKZ_1260. The other recessive candidate genes could not be validated due to technical difficulties (and are likely sequencing artifacts) or due to lack of parental DNA. None of the recessive candidate genes were affected twice or more in this cohort. All predicted deleterious variants were submitted to the ClinVar database https://www.ncbi.nlm.nih.gov/clinvar/ (Landrum et al., 2014).

We inspected the CN profiles from WES-CN and SNP-array for partial overlap with genes affected by heterozygous variant predicted to be deleterious in (recessive) loss of function intolerant or missense intolerant genes (n=48) and could not detect unmasking of a recessive mutation by a CNV. Ultra-rare variants (n=78), X-linked or recessive variants are depicted in Table S4 and uploaded to the ClinVar database (https://www.ncbi.nlm.nih.gov/clinvar/)

### Pathway enrichment analysis of genes affected by rare variants

When looking at the selected protein altering variants (Z-score ≥3, n=44) or loss of function intolerant (PLI ≥0.9, n=4), two relevant pathways were significantly enriched (p-value <1×10^-5^): proliferation and differentiation of smooth muscle cells *(INSR, ITGB1, NOTCH1, TCF4, PDE4D, TERT, ANKRD17, DICER1)* and self-renewal of satellite cells *(ITGB1, NOTCH1).*

### Variant prioritization using different in silico tools

We prioritized all rare variants with three in silico tools (see Methods section). Fifty-four variants in 34 genes had an overlap between VAAST (Yandell et al., 2011, Hu et al., 2013, Kennedy et al., 2014)), which prioritizes based on variant deleteriousness and Phevor and PhenIX which prioritize more on phenotype (Singleton et al., 2014, Zemojtel et al., 2014)). Top ranking variants can be found in Table S3.

Additionally, we found variants in the same gene in multiple patients (Fig. 1). Of these 116 genes (VUS=87, likely deleterious=30), 36 genes were found in ≥3 patients of which six genes were present in more than five patients *(CNTN2, DSPP, NOTCH4, PRRC2A, SEC16B, ZNF717).* Four *(AMBRA1, ATP2A3, DSCAM, NOTCH1)* out of 116 genes were predicted to be intolerant for missense variants (Z-score ≥3). See also Table S4.

**Figure 1.**
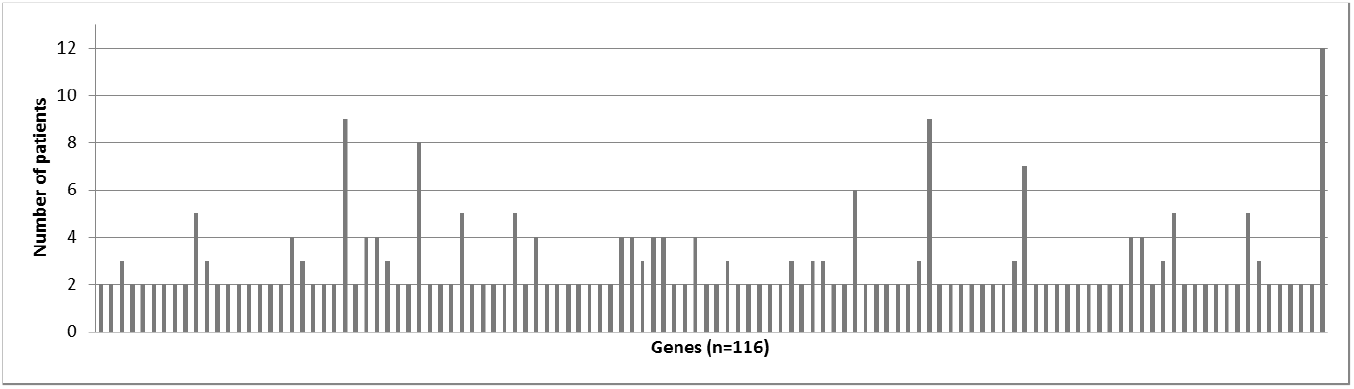
Number of patients with variants per gene. 36 genes were found in >3 patients of which six genes were present in more than five patients *(CNTN2, DSPP, NOTCH4, PRRC2A, SEC16B, ZNF717).* See also Table S6.

### Gene burden analysis

An exome wide gene burden analysis showed ten genes which were enriched for rare putatively deleterious variation compared to the 1000 Genomes project phase 3 samples. The results are shown in Table 3. There were no genes with more than two distinct variants. Each variant was observed only once. A second burden test – only evaluating genes from developmental important pathways and known disease genes – showed no significant difference between our 15 patients and a control group of 44 healthy individuals, who were sequenced in a previous study (Table 4). Also, the number of putative deleterious variants between these two groups was not significantly different (Table 5). Unfortunately, a burden test comparing the variant profiles of these genes between the patients and their parents was not possible since no WES data of the parents was available.

**Table 3.** Results for gene burden analysis

| Gene | Chr. | Start position | rsID | HGVS c | VAAST p-value |
| --- | --- | --- | --- | --- | --- |
| *ALMS1* | chr2 | 73828337 | rs201777220 | c.11885T>C | 1,00E-06 |
| *ALMS1* | chr2 | 73746907 |  | c.9543delA | 1,00E-06 |
| *SLC28A3* | chr9 | 86895771 | rs762396296 | c.1673dupA | 0,002853 |
| *SLC28A3* | chr9 | 86912927 | rs775359011 | c.677G>A | 0,002853 |
| *SP2* | chr17 | 46002296 | rs778698435 | c.1384C>A | 0,002853 |
| *SP2* | chr17 | 45993628 | rs754352270 | c.191C>T | 0,002853 |
| *EPB41* | chr1 | 29435944 |  | c.2410A>G | 0,019802 |
| *EPB41* | chr1 | 29391549 | rs768609152 | c.2063T>G | 0,019802 |
| *AMBRA1* | chr11 | 46563580 | rs755884183 | c.1717C>G | 0,019802 |
| *AMBRA1* | chr11 | 46567274 |  | c.431C>T | 0,019802 |
| *VWA8* | chr13 | 42293773 | rs371770462 | c.3070G>A | 0,019934 |
| *VWA8* | chr13 | 42439870 | rs201163045 | c.1425+2T>C | 0,019934 |
| *CLGN* | chr4 | 141320158 | rs201306926 | c.731A>G | 0,039604 |
| *CLGN* | chr4 | 141315036 | rs200652126 | c.1309G>T | 0,039604 |
| *SDK2* | chr17 | 71357964 | rs147983543 | c.5326G>A | 0,049505 |
| *SDK2* | chr17 | 71415318 |  | c.2173G>A | 0,049505 |
| *PDLIM7* | chr5 | 176911101 | rs200609502 | c.1141A>G | 0,049505 |
| *PDLIM7* | chr5 | 176916502 | rs764108486 | c.761C>T | 0,049505 |
| *GUCY2F* | chrX | 108652306 | rs7883913 | c.1883G>A | 0,019802 |
| *GUCY2F* | chrX | 108638614 | rs35726803 | c.2380G>A | 0,019802 |

N/A = not applicable

**Table 4.** Summary of overlapping top candidate genes

|  | EA/IHPS patients (n=15) |  |  |  | Healthy controls (n=44) |  |  |  |
| --- | --- | --- | --- | --- | --- | --- | --- | --- |
| | Rare (MAF $\leq 0.001\%$) | | Ultra-rare (MAF 0%) | | Rare (MAF $\leq 0.001\%$) | | Ultra-rare (MAF 0%) | |
|  | LB | VUS/LD | LB | VUS/LD | LB | VUS/LD | LB | VUS/LD |
| Important in normal foregut development (Fig. 2) | 1 | 1 | 0 | 0 | 3 | 5 | 1 | 2 |
| Genes associated with genetic syndromes involving both EA and IHPS as variable features (Table 1) | 1 | 0 | 0 | 0 | 2 | 4 | 1 | 1 |
| Genes associated with IHPS (Peeters et al., 2012) | 0 | 0 | 0 | 0 | 0 | 2 | 0 | 0 |
| Genes involved in neuromuscular and connective tissue syndromes associated with IHPS (Peeters et al., 2012) | 1 | 1 | 1 | 0 | 1 | 2 | 1 | 0 |
| Genes involved in syndromes and signaling disturbances associated with IHPS (Peeters et al., 2012) | 0 | 3 | 0 | 0 | 1 | 6 | 1 | 1 |
| Genes involved in ciliopathies and disturbances of gene regulation associated with IHPS (Peeters et al., 2012) | 0 | 1 | 0 | 0 | 4 | 2 | 1 | 0 |
| Genes involved in lymphatic abnormalities and syndromes of environmental and unknown origin associated with IHPS (Peeters et al., 2012) | 0 | 0 | 0 | 0 | 1 | 0 | 0 | 0 |
| Genes involved in genetic syndromes and abnormalities | 1 | 1 | 1 | 1 | 6 | 6 | 0 | 1 |

**Table 5.**
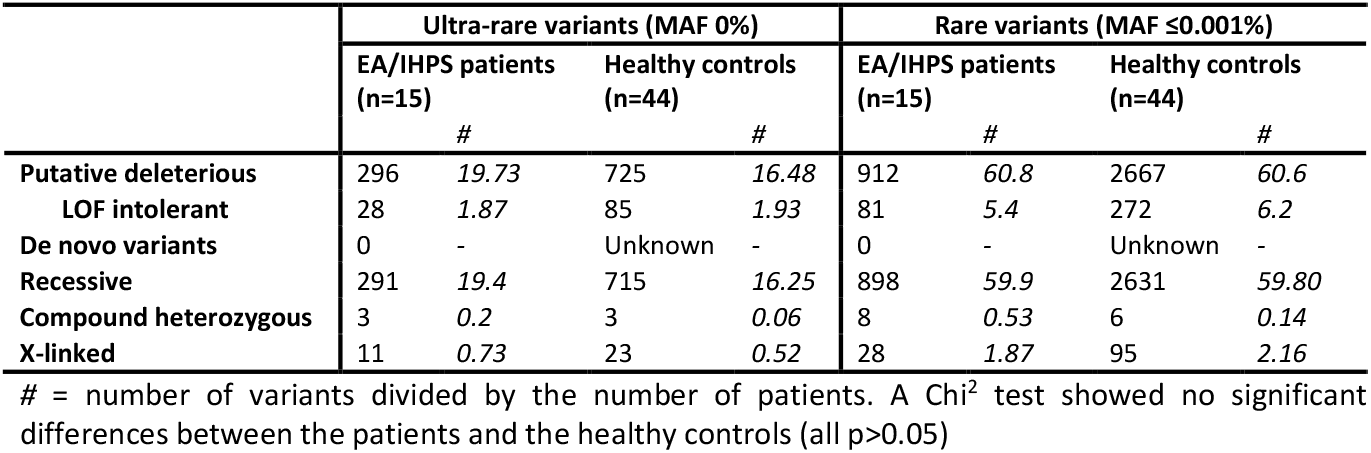
Comparison with control cohort: number of variants

### Expression of main candidate gene during development

With public micro-array transcriptome data we evaluated which genes were upregulated at a specific time-point in the foregut, esophagus or pyloric sphincter and used the output as an indicator of gene expression (see Methods section and Table S5). Of the genes classified as VUS or likely deleterious in our exome sequencing results, 28 genes were upregulated in both the foregut or esophagus as well as the pyloric sphincter: *ADAMTSL4, AGRN, ANKRD29, ARHGAP29, CAMTA1, CDHR5, CNTN2, COL11A1, DNAJC11, HIVEP3, HMCN1, HMGCS2, HSPG2, ITGB3BP, LDB3, MYOF, NKX2-3, NUP133, PCSK9, PKN2, PRDM16, PUM1, RET, SEC16B, SERINC2, TMEM82, VPS13D* and *ZBTB7B.*

Unfortunately, none of the genes enriched in our burden analysis were differentially expressed in mice foregut between E8.5 and E16.5. Seven out of 116 genes with putative deleterious variants in more than one patient were differentially expressed in mice foregut: *Adamtsl4* at E8.5, E14.5 and E16.5; *Ankrd26* at E14.5; *Cntn2* at E8.5, E15.5 and E18.5; *Hspg2* at E8.25, E8.5, E14.5 and E18.5; *Kcnn3* at E8.5 and E15.5; *Ldb3* at E8.5, E14.5 and E15.5; *Sec16b* at E8.5, E14.5 and E16.5. Of the top candidate genes in the manual burden analysis (see Table 4 and Table S6) only *Ret* was differentially expressed in mice at E8.25, E8.5, E11.5, E14.5, E15.5, E16.5 and E18.5.

### Detection of common SNPs associated with IHPS

Determination of the risk allele frequency of four loci highly associated with IHPS (rs11712066, rs573872, rs29784 and rs1933683 near genes *MBNL1, NKX2-5* and *BARX1,* respectively) revealed a significantly higher incidence of rs1933683 in our EA/IHPS cohort compared to the population frequency (OR 3.29 (95% CI 1.27-8.56), p=0.009, see Table S7). The risk allele frequency of the other risk loci was not significantly different from the normal population. We did not detect rare putatively deleterious variants in *MBNL1, NKX2-5* and *BARX1* in the patient exome sequencing data.

## DISCUSSION

We hypothesized that the increased prevalence of IHPS in patients with EA compared to the prevalence of IHPS in the normal population was due to shared CNVs or protein coding alterations in a specific gene, or due to genetic disturbances in genes of shared biological networks during development. As mentioned earlier, both EA and IHPS are variable features in specific genetic syndromes (Table 1). Therefore, to find genetic aberrations that contribute to EA/IHPS we initially searched for pathogenic alterations in known EA or IHPS associated genes (Table S6).

### There are no pathogenic changes in known disease genes

As all parents were unaffected, we started this study by focusing on *de novo,* recessive or X-linked changes affecting these known disease genes. However, we could not identify deleterious protein coding alterations, exonic gains or losses or larger CNVs affecting these genes. This is in line with previous studies in which limited causal changes could be detected in patients with EA and associated anomalies (Zhang et al., 2017, Hilger et al., 2015, Brosens et al., 2016b)

Next, we extended our analysis to all genes covered in the exome capture. Given the small sample size (n=15), the low prevalence of the disorder and the high impact on development, we concentrated on genes intolerant to variation (Lek et al., 2016, Ruderfer et al., 2016) harboring rare putative deleterious single nucleotide changes or large CNVs. Moreover, we determined the segregation of alterations in these candidate genes, putative recessive (X-linked, compound heterozygous and homozygous recessive) and all ultra-rare protein altering changes absent from the gnomAD database (Lek et al., 2016) as the later have a high chance of being *de novo* (Bennett et al., 2017). Unfortunately, we did not identify any *de novo* mutations or *de novo* CNVs. None of the identified inherited rare CNVs overlapped in these patients. We could confirm the presence of a compound heterozygous variant in *FAM46A* in one patient and an X-linked variant in *SH3KBP1* in another patient. However, *FAM46A* and *SH3KBP1* are not known to be associated with the gastrointestinal or respiratory tract and were not differentially expressed at the time points important for foregut morphogenesis. These findings made us conclude that neither a dominant nor a recessive model can explain the combination of EA and IHPS in these patients.

### The coding sequences of genes crucial in esophageal and pyloric sphincter formation are affected

Subsequently, we focused on genes involved in foregut development. Literature research together with data of previous expression studies provided an overview of genes important for foregut development (Fig. 2).

**Figure 2.**
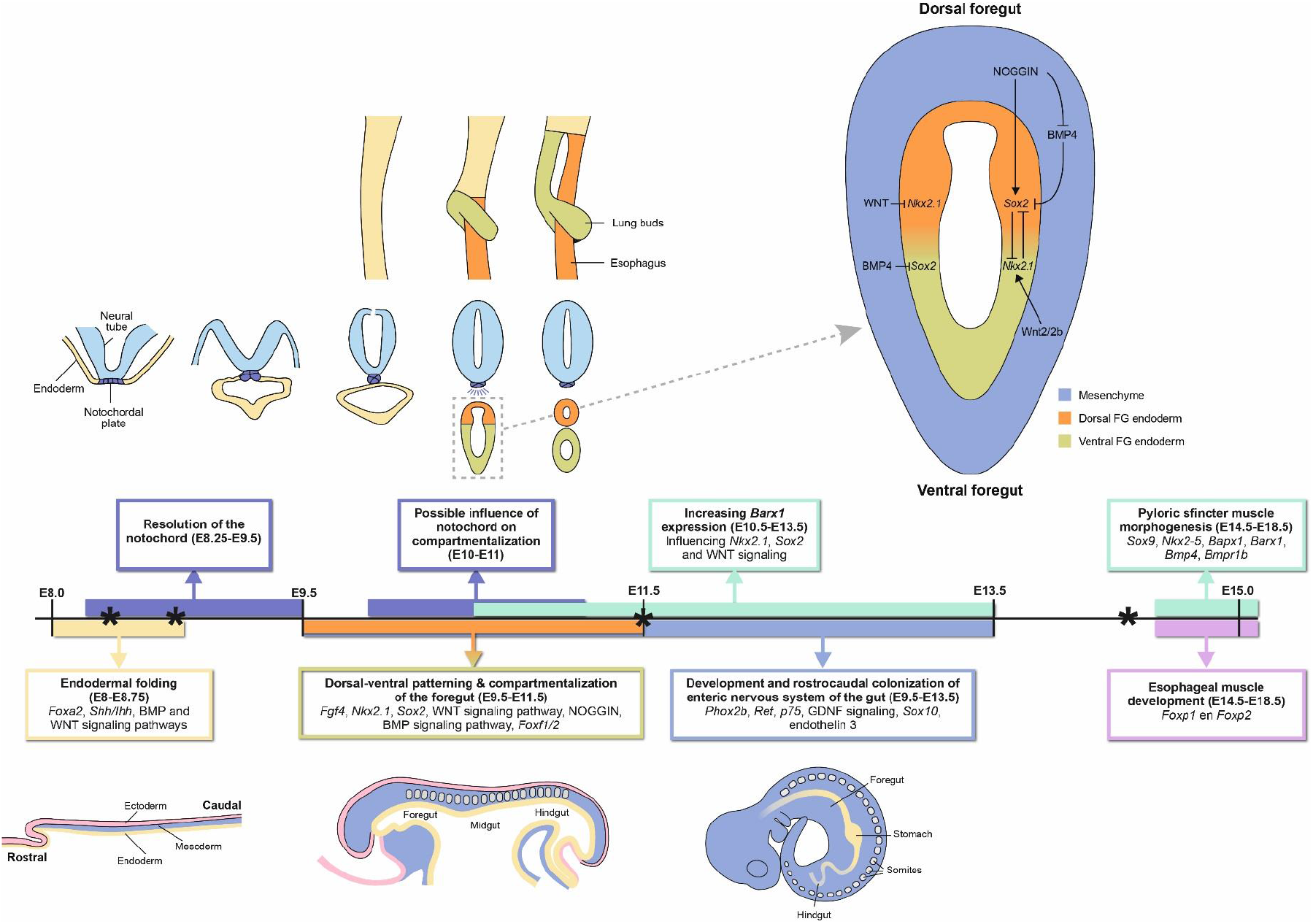
Timeline of models and genes known to be important for foregut development in mice. (Heath, 2010, Fausett and Klingensmith, 2012, Perin et al., 2017, Anderson et al., 2006), * = time points used in expression analysis (see Table S5)

The development of the foregut is most studied in mouse models. In mice, early foregut formation starts with *Foxa2* stimulation of the anterior endoderm at E8.0 (Heath, 2010). The endodermal sheet folds and forms a tube at E8.75 (Sherwood et al., 2009). Next, signals from the notochord start dorsal-ventral patterning around E9.0, with high Nkx2.1/absent *Sox2* in the ventral future trachea and absent Nkx2.1/high *Sox2* in the dorsal future esophagus and stomach (Que et al., 2007). These dorsal-ventral patterns lead to compartmentalization of the foregut. Between E9.5 and E11.5 the foregut separates in the primordial esophagus and stomach, and in the primordial trachea. Primordial lung buds become apparent at E9.5 (Sherwood et al., 2009). The separation site is marked by mesenchymal expression of *Barx1* (Woo et al., 2011). The esophagus is completely separated from the trachea at E11.5.

Pyloric sphincter formation is mostly studied in chick and mouse models. This formation starts with the thickening of the circular smooth muscle layer between the antrum and the duodenum around E14.5 and the primordial pyloric sphincter is complete around E18.5 (Smith et al., 2000a, Self et al., 2009). In addition to its functioning in foregut separation, the *Barx1* homeobox gene is also vital for stomach differentiation and stomach smooth muscle development. It inhibits *Wnt* signaling (Woo et al., 2011) and modulates the expression of *Bapx1,* another important factor required for pyloric sphincter morphogenesis (Jayewickreme and Shivdasani, 2015, Stringer et al., 2008, Verzi et al., 2009).

Given the described importance of these genes in normal development, we hypothesized that variations in multiple genes important for foregut morphogenesis might explain the higher incidence of IHPS in patients with EA. We compared a selection of genes – known to be important for foregut morphogenesis or syndromatically associated with EA or IHPS – between the patients and the healthy controls (Table 4, Table S6). Interestingly, in *TNXB* (NM_019105.6:c.4444G>A, p.Val1482Met), *WDR11* (NM_018117.11:c.1138G>T, p.Val380Phe), *PEX3* (NM_003630.2:c.1012A>G, p.Ser338Gly), *TBX3* (NM_016569.3:c.506G>A, p.Arg169Gln), and *GDF6* (NM_001001557.2:c.281C>G, p.Pro94Arg) rare variants were present that were absent in the control group. These variants might not be sufficient to result in disease but are predicted to impact the protein and might contribute together with other unknown factors to disease development.

### There is no high frequency burden of rare variants

Given the limited number of samples, we will only detect a gene burden if it is large and has a high impact. We compared the total rare and ultra-rare variant burden of putative deleterious variants in all genes. The number of ultra-rare variants was slightly higher in the patient group compared to the control group but did not differ significantly (Table 5). A second burden analysis identified ten genes with more variants compared to those seen in the 1000 Genome cohort, two variants were predicted to be deleterious (Table 3). Unfortunately, they did not show any overlap with the results of the expression analysis or candidate genes selected from the literature. Therefore, these variants are not likely to explain the increased incidence of IHPS in EA patients or EA/IHPS development. A rare variant burden might exist but we could not detect it due to limited sample size and/or focus on known candidate genes.

Of all the protein coding changes classified as VUS or higher (Table S4; Table S8), 116 genes were affected with a variant in more than one patient (Fig. 1). Seven of these genes *(ADAMTSL4, ANKRD26, CNTN2, HSPG2, KCNN3, LDB3, SEC16B)* were differentially expressed in the developing foregut, esophagus or pyloric sphincter in mice between E8.25 and E16.5. However, none of these genes could explain the combination of EA and IHPS within a patient based on their function; none of these genes is known to be associated with the gastrointestinal or respiratory tract. Furthermore, most variants had a population frequency above the prevalence of EA/TEF. If these variants are highly penetrant, they would not be the likely cause. Increasing sample sizes (drastically) would allow an analysis going beyond known intolerant genes, allow us to consider reduced penetrance and potentially identify a shared genetic etiology.

### Known common variants associated with IHPS development could have an impact in some patients

Since certain SNPs have been identified with GWAS to be highly associated with IHPS, we wondered if these known common haplotypes could also play a role in the higher incidence of IHPS in patients with EA. In our cohort, we found a significantly higher incidence of the risk loci rs1933683 compared to the population frequency (Table S7). Three patients were homozygous for the risk allele and have a substantially increased risk for IHPS development. The common risk haplotype might therefore impact IHPS development in some of the IHPS patients. However, further research is needed to confirm the impact of this haplotype in a larger EA and EA/IHPS population.

### Possible contribution of non-genetic factors

All the data presented so far made us conclude that dominant *de novo* variations in possible disease causing genes do not play a role in our cohort. Recessive inheritance cannot totally be excluded, although our results are not suggestive for this mode of inheritance. We did identify in all patients putative disease-causing variants. Nevertheless, as all parents from whom these variants were inherited were not affected, these variants could contribute but not cause the disease. Previous studies suggested the contribution of non-genetic factors as an explanation for the combined occurrence of EA and IHPS.

### Could IHPS be an acquired condition related to surgery or treatment of EA?

The overrepresentation of IHPS in EA patients made us wonder if IHSP could also be the result of the atresia itself, potentially as a result of the surgical procedure to correct the atresia or the result of treatment. Previous studies also mentioned vagal nerve lesions, a gastrostomy and transpyloric feeding tubes as possible causes for an increased incidence of IHPS after correction of EA (Ilhan et al., 2018). IHPS has been suggested to be a neuromuscular disorder with the involvement of smooth muscle cells, interstitial cells of Cajal and the enteric nervous system. The hypertrophy is suggested to be the result of discoordinated movements of the pyloric sphincter and the contractions of the stomach (Hayes and Goldenberg, 1957), perhaps as the result of absent nitric oxide synthase activity (Vanderwinden et al., 1992). Impaired gastric contractility and esophageal relaxation were observed in Adriamycin and doxorubicin induced EA in mice (Tugay et al., 2003, Tugay et al., 2001). Mechanistically, this association between EA and IHPS seems plausible. However, it does not explain why IHPS is not fully penetrant in patients with EA. The most common thought is that mechanical and environmental factors disturb the developmental field. To which extent these factors influence the development of the child, depends on the specific risk factors and their timing. Further research on the cause and other specific clinical risk factors for patients with EA should be considered, e.g. the late start of oral feeding or the long-term feeding through a tube instead of drinking themselves.

### Models for EA/IHPS disease etiology

Since we hypothesized that genetic defects, disturbing foregut morphogenesis, would be responsible for the combination of EA/IHPS, we started with the thought of a (monogenetic) syndromic model. However, we have not been able to find a central gene impacted in most patients which can explain the increased prevalence of IHPS in patients with EA. An as yet unknown syndrome is unlikely since we have not found *de novo* (as all parents are unaffected) or shared high impact variants in the same gene multiple patients. However, we cannot exclude *de novo* mutations which have been seen in the GnomAD exomes or genomes, nor did we look beyond the coding part of our genome.

Furthermore, we have detected inherited rare variants in candidate genes and genes affected more than once by variants with a low (unaffected) population frequency. Therefore, we cannot exclude a genetic component. Another option is that, although the combination of EA and IHPS could not be explained by one gene or locus, the underlying cause of EA and the other associated anomalies is the result of hits in multiple genes. About 10% of the patients with EA have an underlying genetic syndrome (Brosens et al., 2014) and more can be expected. The phenotypical spectrum of this cohort is very heterogeneous and could be the result of impacts on multiple genes. IHPS could then independently be caused by mechanical factors such as the surgical procedure.

Furthermore, environmental risk factors have been suggested for EA and IHPS, like pesticides, smoking, herbicides and periconceptional alcohol or multivitamin use (Zwink et al., 2016, Felix et al., 2008, Feng et al., 2016, Markel et al., 2015, Krogh et al., 2012, Sorensen et al., 2002). Considering the absence of highly penetrant recurring genetic variations, we now hypothesize different multifactorial models for disease development. In all these models the combination of CNVs, deleterious protein alterations (Felix et al., 2007b, Brosens et al., 2014), severe changes in the developmental field during the organogenesis (Martinez-Frias, 1994, Martinez-Frias and Frias, 1997) and/or environmental inducing epigenetic changes (Sorensen et al., 2002) together can modulate the phenotypical spectrum seen in these patients. Examples of possible mechanical and environmental factors disturbing the developmental field are mice models with Adriamycin induction or dorsal-ventral patterning signals from the notochord.

Our first hypothesis is based on the earlier published theory of Brosens et al. about disturbed ENS development (Brosens et al., 2016a). It includes a seesaw model in which risk factors are in balance with protective mechanisms. In this model, the fulcrum can be shifted by a genetic variation in a central gene, which automatically disrupts the balance. When applying this theory on EA/IHPS patients, this would lead to more or less affected organ systems within the VACTERL syndrome. However, in this study we have not been able to detect one central gene. This makes a seesaw model with a variable fulcrum less likely as an explanation for the increased prevalence of IHPS in the EA population.

A second hypothesis is a burden model (Fig. 3A). Similar, (epi)genetic, environmental and mechanical factors form a burden of risk factors, which balances with protective mechanisms. In this model, the point of balance is not shifted by a mutation in a central gene and every person has contributions of certain risk factors. But in most cases this does not lead to affected organ systems. There is an intermediate range between normal and affected in which individuals can have the genetic burden but lacks an abnormal phenotype (reduced penetrance) or their symptoms differ in severity (variable expressivity). The latter would fit the results in this study; maybe we did have detected variants but have we failed to interpret them correctly as parents were seemingly unaffected and/or the variant frequency can be higher in unaffected controls to be of relevance in patients. Mechanical or environmental factors could have made the difference in shifting the balance. All together the burden model is a plausible explanation for the disease development.

**Figure 3.**
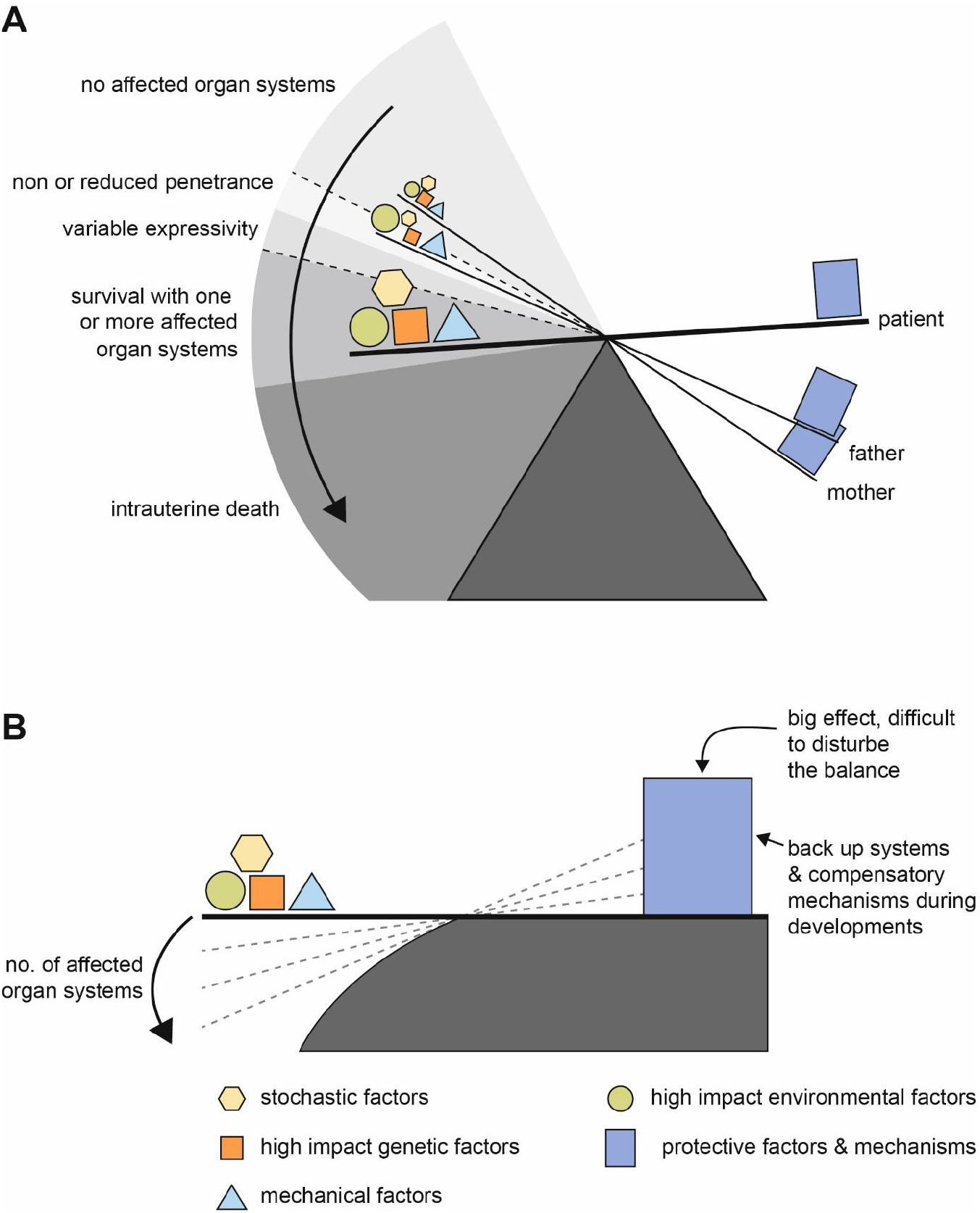
Two models for EA/IHPS etiology A = burden model, B = slippery slope model. The combination of multiple high impact factors (genetic, environmental, mechanical and/or stochastic) together can modulate the phenotypical spectrum. These risk factors are in balance with protective factors like backup systems and compensatory mechanisms.

Last, we hypothesize a slippery slope model (Fig. 3B). In this model, the burden of low impact genetic variants and environmental disturbances alone does not impact the balance seen in the seesaw model unless it crosses a certain threshold. Moreover, we hypothesize that the protective mechanisms (e.g. compensatory mechanisms) during development are very strong, making it really difficult to shift the balance. Most fetuses do not develop any malformations despite the combined genetic and environmental burden or do not survive. But once the threshold is reached, the balance is immediately greatly disrupted and often multiple organ systems are affected. This model fits with the phenotypical results in this study since four patients (14.8%) had three or more anomalies within the VACTERL spectrum. In this model there is a high tolerance for low impact genetic variation and only high impact variation (aneuploidies, exposure to toxic substances, pathogenic changes in developmental crucial genes) shifts the balance. When the balance is disturbed, it shifts drastically. We did not detect high impact changes responsible for the EA/IHPS combination. As parents are unaffected it is (in this model) unlikely that inherited variants impact disease development, nor would variants which are seen in the (unaffected) population controls.

### Limitations

Not finding any positive correlation between DNA variations in specific genes or developmental pathways is partly due to the small data set we have (15 patients). The small sample size is no problem for the *de novo* and our recessive model strategy in known disease genes, but it is so for the heterozygous variant burden analysis. Another limitation is the lack of data on the expression of genes involved in normal foregut development in human embryos. Our gene selection was based on mouse transcriptome data. Little human data is available since human embryos of 4 to 6 weeks old are generally not preserved. However, although it is unclear how precisely the foregut development in mice corresponds with humans, it is unlikely that this is very different in its early phases. Finally, one could argue that variations in the non-coding part of the genome are major contributors. Although we did investigate known IHPS risk loci and determined genome wide CNV profiles, we did not determine genome wide variation.

### Conclusions

To conclude, *de novo* mutations (a dominant model) and homozygous or compound heterozygous mutations (a recessive model) in the protein coding part of the genome are not a likely cause for the combination of EA and IHPS. Although the presence of genetic variation in likely candidate genes suggests a genetic component, there does not seem to be an enrichment of genetic variants in good candidate genes in patients. There are putative deleterious variants in foregut or disease genes which might contribute to disease development and although there is no difference in burden, some variants might contribute more than others and this is not taken into account in a burden test. We might have misinterpreted the impact of some of the inherited variants. Furthermore, in some patients the IHPS predisposing locus rs1933683 is present.

We hypothesized several multifactorial models in which the combination of multiple high impact genetic, mechanical and environmental factors together can shift the balance from normal to abnormal development. A burden model with reduced penetrance or variable expressivity is most likely if genetic factors contribute. Future research should investigate the incidence of IHPS in bigger EA patients cohorts to further explore this theory. To exclude the role of treatment or surgery, clinical factors related to the surgical correction of EA – for example vagal nerve lesions after surgery, the late start of oral feeding or transpyloric feeding tubes – should be systematically registered.

## MATERIAL AND METHODS

### Patient cohort

This study was approved by the Medical Ethical Review Board of Erasmus MC – Sophia Children’s Hospital (MEC 193.948/2000/159). We searched the Erasmus University MC-Sophia EA-cohort and the database of the standardized prospective longitudinal follow up program in our hospital for children with congenital anatomical anomalies (Gischler et al., 2009) for patients born between 1970-2017 with a combination of both EA and IHPS in history. Patients were included and analyzed after parental informed consent

### SNP-array analysis

Micro-array analysis was performed using the single-nucleotide polymorphism (SNP) CytoSNP-850Kv0 BeadChip (Illumina Inc., San Diego) using standard protocols and the GenomeStudio genotyping module (v1.9.4, www.illumnia.com). Visualization of Copy Number Variations (CNVs), Runs of Homozygosity (ROH) and comparisons to in-house control cohorts as well as published cohorts of affected and control individuals was done using Biodiscovery Nexus CN7.5. (Biodiscovery Inc., Hawthorne, CA, USA) and described previously (Brosens et al., 2016b).

### Variant pre-filtering and prioritization

The initial variant filtering method has been described previously (Halim et al., 2017). In brief, we included all variants with an allele frequency below 1% in 1000 Genomes phase 3 version 5, Exome Variant Server 6500 v0.0.30, Genome of the Netherlands (Genome of the Netherlands, 2014), ExAC 0.3 and our in-house cohort (n=906), consisting of individuals captured with the SureSelect Human All Exon 50 Mb Targeted exome enrichment kit v4 (n=279), SureSelect Clinical Research Exome v1 (n=387) and Haloplex Exome target enrichment system (n=240), Agilent Technologies, Inc., Santa Clara, California).

All nonsense variants, variants predicted to affect splicing and all variants with a Combined Annotation-Dependent Depletion (CADD) score (Kircher et al., 2014) above 20 were selected for individual patient analysis in downstream tools. Different downstream tools were used to prioritize the variants. Prioritized variants were further classified according to the criteria in Table S8. Determination of variant segregation and confirmation of *de novo* of inherited status of variants was done with Sanger sequencing unless otherwise indicated.

### Variant burden test and prioritization using Opal

We used the Variant Annotation, Analysis & Search Tool (VAAST) (Yandell et al., 2011, Hu et al., 2013, Kennedy et al., 2014) cohort analysis embedded in Opal 4.29.5 (Fabric Genomics, Oakland, CA, USA) to rank the variants in the individual patients. Secondly, we performed a burden test on the full exomes using Exome Variant Server 6500 v0.0.30 and 1000 Genomes phase 3 version 5 as a control cohort. We used a 1% allele frequency cut-off for recessive (hemizygous and homozygous) variants and 0.1% cut-off for heterozygous variants. Compound heterozygosity was not considered in this analysis as we did not know the phase of the haplotypes. Only putative protein changing (nonsense, missense, initiator codon variants, in-frame indels, splice sites and splice regions) variants were taken into account. Since we were only interested in putative deleterious variants we used an Omicia score of 0.79 as a threshold as this cut-off has a false positive rate of 5%. Omicia is an algorithm included in the Opal software that combines SIFT (Ng and Henikoff, 2001), PolyPhen (Adzhubei et al., 2010), MutationTaster (Schwarz et al., 2010) and PhyloP (Siepel et al., 2005) to predict deleteriousness of variants.

For the VAAST burden test we used a minimum significance of 0.05 and a gene had to have at least two distinct variants in the case set. These genes were used as a gene panel in the individual patient analysis. Individual variants were prioritized before individual inspection as follows. First, all recessive (X-linked and putative homozygous and compound heterozygous), putative rare (MAF ≤0.001%) and damaging *de novo* variants were selected. Secondly, the top 10 of variants ranked by the VAAST 1.1 prioritization algorithm and subsequently the top 10 variants re-ranked by the Phevor algorithm (Singleton et al., 2014) were included. We used the Human Phenotype Ontology (HPO) (Singleton et al., 2014) terms esophageal atresia and pyloric stenosis as phenotype terms in the algorithm. Finally, variants passing the pre-filtering criteria in genes from the burden test were included.

### Variant prioritization using bioinformatic genotype-phenotype correlation tools

Three modules were used: PhenIX (Zemojtel et al., 2014) (http://compbio.charite.de/PhenIX/), the Exomiser (Robinson et al., 2014) (http://www.sanger.ac.uk/resources/software/exomiser/submit/) and the HPO prioritization incorporated within the Cartagenia software. Settings were as followed.

Using PhenIX the full patient phenotype in HPO terms was used, the exome target region filter is on and allele frequency filter of 0.1%, pathogenicity filter was on and mode of inheritance unknown. Genes were prioritized using PhenIX which compares patient phenotypes against human phenotypes only. As a cut-of we used a gene relevance score of 0.8 in combination with a variant score of 0.8, or a total score of 0.9.

When using the Exomiser tool we used similar settings: full patient phenotype in HPO terms, exome target region filter is off, allele frequency filter 0.1%, pathogenicity filter on. We did not remove dbSNP variants nor used an inheritance model. Genes are now prioritized using hiPhive, which compares phenotypes against all species. As a cut-of we used a phenotype score of 0.8 in combination with a variant score of 0.8, or an Exomiser score of 0.9.

### Pathway enrichment analysis of genes affected by rare variants

To investigate if specific pathways are enriched with ultra-rare variants, Gene IDs with variants in canonical splice sites (n=16), nonsense variants (n=21), protein altering inframe InDels (n=28) and missense variants (n=557) were uploaded to Ingenuity pathway Analysis (Qiagen, Venlo, The Netherlands). Additionally, a more stringent set was uploaded with loss of function variants, predicted to be loss of function intolerant (PLI ≥0.9, n=4) and protein altering variants with a Z-score ≥3 (n=44).

### Expression of candidate genes

Candidate gene expression was determined at relevant developmental time points in human and mouse. Gene expression of top-ranking genes derived from the burden analysis and individual patient sample prioritizations were determined using datasets (GSE13040, GSE19873, GSE34278, GSE15872, GSE43381) downloaded from the Gene Expression Omnibus (GEO) (Edgar et al., 2002). We used public data on mice on the endoderm, mesoderm and ectoderm at E8.25, foregut at E8.5 and esophagus, stomach, pyloric sphincter and intestine at E11.5-E18.5 (https://www.ncbi.nlm.nih.gov/geo/) (Stephens et al., 2013, Li et al., 2009, Sherwood et al., 2009, Millien et al., 2008, Chen et al., 2012). These datasets were imported into BRB-ArrayTools Version: 4.5.0 – Beta_2. (http://linus.nci.nih.gov/BRB-ArrayTools.html), annotated by Bioconductor (www.bioconductor.org), R version 3.2.2 Patched (2015-09-12 r69372) and normalized. We determined differential expression between tissue types and classified upregulated genes being expressed in the tissue under investigation.

### Detection of common SNP associated with IHPS

Genome-wide association studies (GWAS) revealed five loci highly associated with IHPS (rs11712066, rs573872, rs29784, rs1933683 and rs6736913), pointing towards *MBNL1, NKX2-5, BARX1* and *EML4* as candidate genes (Feenstra et al., 2012, Everett and Chung, 2013, Fadista et al., 2019). Since rs6736913 is a low frequency missense variant, we did not further analyze this SNP in our patients. We used Sanger sequencing to determine the risk allele frequency of the other four SNPs. With a chi square test we compared the allele frequency in our patients with the gnomAD dataset (http://gnomad.broadinstitute.org/).

## ACKNOWLEDGEMENTS

We are grateful for the help of families, patients and the cooperation of the patient cooperation “Vereniging voor Ouderen en Kinderen met een Slokdarmafsluiting”. We would like to thank Tom de Vries Lentsch for preparing the figures.

## COMPETING INTERESTS

Authors do not have any potential conflicts (financial, professional, or personal) relevant to the manuscript to disclose.

## FUNDING

Erwin Brosens was funded by the Sophia Foundations for Scientific Research, projects SSWO-493 and SWOO13-09.

## DATA AVAILABILITY

Variants included in Table S4 are submitted to the ClinVar database. (http://www.ncbi.nlm.nih.gov/clinvar/)

## AUTHOR CONTRIBUTIONS STATEMENT

Conceptualization: D.T., R.W., R.H., A.K., E.B.; Methodology: R.B., H.E., W.IJ., A.K., E.B.; Software: R.B., T.B.; Validation: C.K., V.M., D.H.; Formal analysis: C.K.; Investigation: C.K., Resources: Y.B., N.B., A.B., H.E., H.IJ.; Data curation: E.B.; Writing – original draft: C.K., R.H., E.B.; Writing – review & editing: C.K., R.B., Y.B., A.B., H.E., W.IJ., H.IJ. D.T., R.W., R.H., A.K., E.B.; Visualization: C.K., Supervision: E.B.; Project administration: D.T., R.W., R.H., A.K., E.B.; Funding acquisation: D.T., R.W., R.H., A.K.

